# Chimeric RNA:DNA Donorguide Improves HDR *in vitro* and *in vivo*

**DOI:** 10.1101/2021.05.28.446234

**Authors:** Brandon W. Simone, Han B. Lee, Camden L. Daby, Santiago Restrepo-Castillo, Gabriel Martínez-Gálvez, Hirotaka Ata, William A.C. Gendron, Karl J. Clark, Stephen C. Ekker

**Affiliations:** Department of Biochemistry and Molecular Biology, Mayo Clinic, Rochester, MN; Mayo Clinic Graduate School of Biomedical Sciences, Virology and Gene Therapy Track, Mayo Clinic, Rochester, MN; Mayo Clinic Graduate School of Biomedical Sciences, Biomedical Engineering and Physiology Track, Mayo Clinic, Rochester, MN; Department of Clinical and Translational Sciences, Mayo Clinic, Rochester, MN

**Author notes:** **Corresponding author:** Stephen C. Ekker-, Department of Biochemistry and Molecular Biology, Mayo Clinic, 200 1st 10 Street SW, Rochester, MN 55905.

## Abstract

Introducing small genetic changes to study specific mutations or reverting clinical mutations to wild type has been an area of interest in precision genomics for several years. In fact, it has been found that nearly 90% of all human pathogenic mutations are caused by small genetic variations, and the methods to efficiently and precisely correct these errors are critically important. One common way to make these small DNA changes is to provide a single stranded DNA (ssDNA) donor containing the desired alteration together with a targeted double-strand break (DSB) at the genomic target. The donor is typically flanked by regions of homology that are often ~30-100bp in length to leverage the homology directed repair (HDR) pathway. Coupling a ssDNA donor with a CRISPR-Cas9 to produce a targeted DSB is one of the most streamlined approaches to introduce small changes. However, in many cell types this approach results in a low rate of incorporation of the desired alteration and has undesired imprecise repair at the 5’ or 3’ junction due to artifacts of the DNA repair process. We herein report a technology that couples the spatial temporal localization of an ssDNA repair template and leverages the nucleic acid components of the CRISPR-Cas9 system. We show that by direct fusion of an ssDNA template to the trans activating RNA (tracrRNA) to generate an RNA-DNA chimera, termed Donorguide, we recover precise integration of genetic alterations, with both increased integration rates and decreased imprecision at the 5’ or 3’ junctions relative to an ssODN donor *in vitro* in HEK293T cells as well as *in vivo* in zebrafish. Further, we show that this technology can be used to enhance gene conversion with other gene editing tools such as TALENs.

## Introduction

Targeted genomic alterations have revolutionized the field of molecular biology. Most recently, these efforts have been focused on designer nucleases such as the clustered regular interspersed short palindromic repeat (CRISPR) and their corresponding CRISPR associated proteins (Cas), namely Cas9 ^1^ ^2^ ^3^. These Cas proteins can be directed to a specific region in the genome specified by an ~20nt guide RNA and induce a targeted double-strand break ^3^. In the absence of a repair template, the endogenous DNA repair machinery will most likely repair the lesion by re-ligating the free DNA ends by an un-templated repair pathway such as non-homologous end joining (NHEJ) ^4^ or alternative end joining (Alt-EJ) including microhomology-mediated end joining (MMEJ) ^5^ ^6^. In the presence of a repair template such as a single stranded oligo deoxy nucleotide (ssODN) or a double stranded DNA (dsDNA) template in the form of a PCR product or plasmid, precise repair is triggered that is directed by regions of homology on the donor template that mimic endogenous DNA repair using a chromosome as a repair template ^7^. Gene replacement via dsDNA template has been shown to be RAD-51 dependent ^8^ ^9^ and be active in G2/M of the cell cycle when a homologous chromosomal template would be present ^10^. Conversely, single stranded- template repair (SSTR) has been shown to be RAD-51 independent ^11^ and be inactive during G2/M ^12^. Further, SSTR has recently been found to be dependent on the Fanconi Anemia (FA) pathway ^7^.

Repair with a dsDNA donor is relatively inefficient in most cell types ^13^. Alternatively, repair via an ssDNA donor has been shown to be efficient for small genetic changes in multiple cell types ^14^ ^13^ ^15^. Several studies have demonstrated that ssODNs are sufficient donors for inserting small (<50bp) changes into the genome with appreciable rates ^16^. Although many gene replacement strategies with dsDNA use long homology arms of 0.5kbp-1kbp, ssDNA-mediated gene replacement can proceed with homologies much shorter than this ^17^. In fact, the length of flanking homologies has been shown to be decreasingly important with the insertion of smaller cargos into the genome ^18^ ^17^. For insertion of smaller cargos such as single base pair insertions or substitutions, single stranded oligos with at least 30bp of flanking homology are typically employed.

Though it is a relatively efficient process *in vitro*, rates of homology directed repair (HDR) with Cas9-mediated double strand breaks (DSBs) coupled with an ssODN repair template in zebrafish are typically quite modest ^19^ ^20^ and often result in imprecise repair at either the 5’ or 3’ junction that is thought to be an artifact of the DNA repair process ^21^. In zebrafish, the imprecision at the insertion junctions has been explained, at least partly, by the stochastic integration of fragments of the ssODN repair template ^22^.Though ssODN template induced changes are typically sufficient for studying a clinically relevant mutation or generating a transgenic cell line, the low efficiency and imprecise nature of the insertion events can render this methodology difficult to deploy in applications that require single base-pair precision.

Recent work has shown that the spatial-temporal localization of a repair template via covalent linkage of an ssDNA donor to the Cas9 nuclease increases rates of HDR significantly ^23^ ^24^. However, these approaches require complex protein-nucleic acid engineering for each target locus and therefore preclude them from being widely adapted approaches for precision gene engineering. These observations using a protein-nucleic acid fusion indicate that methods that improve spatial temporal location of the repair template increase rates of gene replacement.

Several methods have been developed to preclude the need for the traditional targeted DSB coupled with a homologous repair template, including base editing (BE) ^25^, ^26^ and prime editing (PE) ^27^. Though these technologies do not require a DSB followed by HDR with a DNA repair template, they are not without their drawbacks. Even with several iterations of the BE system, the editing window remains relatively large and makes high throughput and precision applications difficult. Additionally, prime editing requires several large and potentially toxic components to be delivered to the system and relies on successful delivery and activity of a viral reverse transcriptase.

Here we report a methodology that is backwards-compatible with common CRISPR technologies available to most research laboratories that uses the spatial-temporal components of the CRISPR-Cas9 system and small flanking homologies sufficient for small DNA modifications. We demonstrate that by direct covalent fusion of an ssDNA repair template to the tracrRNA, deliverable as CRISPR-Cas9 nuclease RNP results in precise gene conversion with as little as 10bp of homology when coupled with a CRISPR-Cas9 mediated DSB. We demonstrate that our approach is amenable both *in vitro* as well as *in vivo* using human embryonic kidney (HEK293T) cells and zebrafish, respectively. Additionally, we find that Donorguide can enhance the activity of other gene editing tools such as TALENs to increase gene modification. Taken together this work provides a streamlined approach that requires only a single RNP complex be delivered to induce precise molecular alterations at rates comparable to the imprecise repair of ssODN-mediated HDR.

## Results

### Donorguide SpCas9 introduces precise insertions *in vitro*

To assess the feasibility of covalently tethering a DNA template to the 3’ end of a tracrRNA to induce precise genetic edits, we employed the HEK 293T cell system due to their ease of transfection (Figure 1). We previously identified a highly active *Streptococcus pyogenes* Cas9 (SpCas9) target site in the safe harbor region AAVS1 ^28^ (Supplemental Figure 1) and determined this to be a candidate locus for our approach to ensure that SpCas9 was active at the target locus. Due to synthetic RNA synthesis limits, we used the separate CRISPR RNA (crRNA) tracrRNA system instead of the single guide RNA (sgRNA) system. We fused a DNA repair template to the 3’ end of the tracrRNA containing 12bp of homology flanking a 2bp insertion (Fig 2a) generating a chimeric RNA:DNA hybrid. This chimeric tracrRNA was annealed with a canonical SpCas9 crRNA to form what we term the Donorguide and combined with SpCas9 recombinant protein to form the active RNP complex. HEK 293T cells were electroporated with the SpCas9 Donorguide RNP and screened for genetic insertions 48 hours later by next generation sequencing (Fig 1). We found that the Donorguide SpCas9 RNP introduced the precise 2bp insertion at ~2% efficiency (Fig 2B). In addition, we found that Donorguide introduced indel mutations at markedly lower rates than a canonical tracrRNA. These data suggested that the Donorguide methodology is a viable approach to introduce precise small molecular changes *in vitro*, albeit at modest rates.

**Figure 1.**
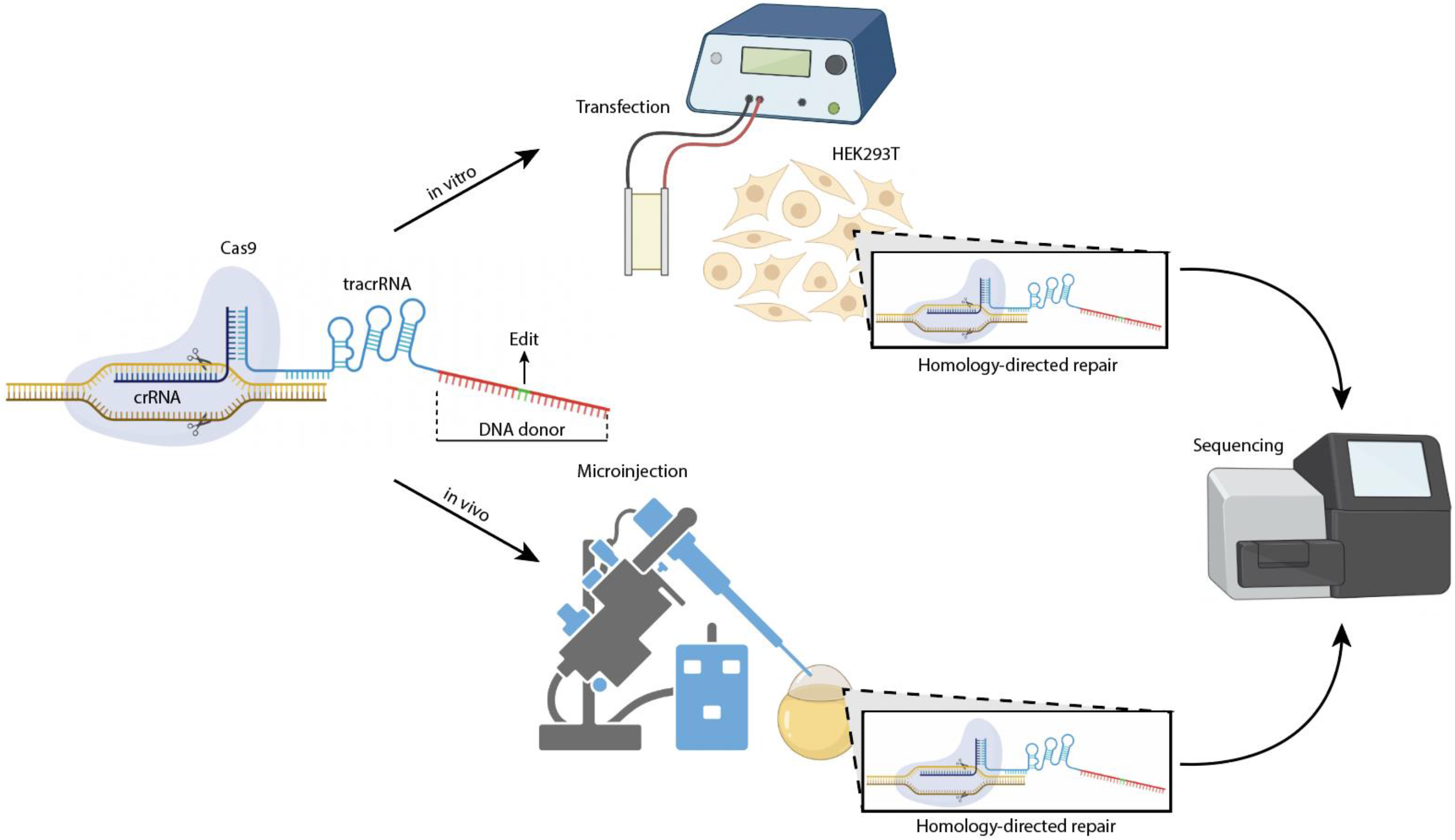
Donorguide composition and experimental workflow. (Left) Donorguide consists of a canonical CRISPR RNA (crRNA), trans activating crRNA (tracrRNA) that is fused to a DNA repair template (Top) *In vitro* experiments were carried out by electroporating Donorguide RNP complexes into cells followed by next generation sequencing (Bottom) *In vivo* experiments were carried out by microinjecting single cell zebrafish embryos with Donorguide RNP complexes cells followed by next generation sequencing

**Figure 2.**
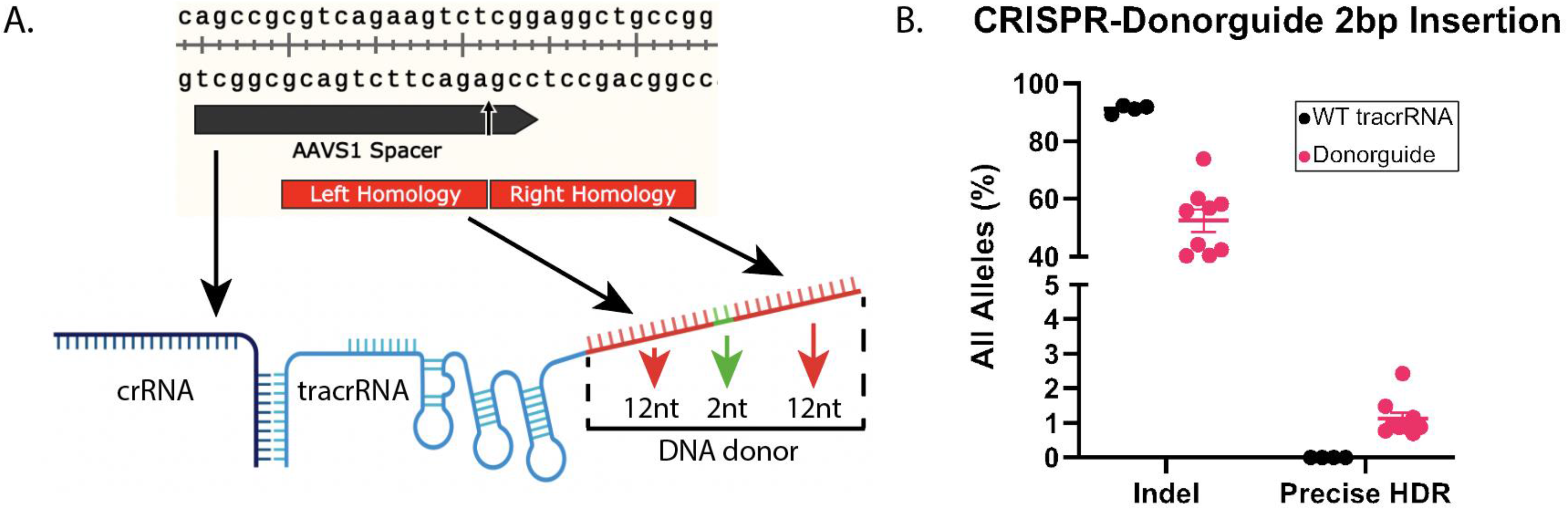
*In vitro* validation of CRISPR Donorguide at AAVS1. **(A)** AAVS1 CRISPR Cas9 target site and corresponding CRISPR Donorguide Design. A 2nt insert was introduced flanked by 12nt of homolgy to the genomic target site. **(B)** Quantification of CRISPR Donorguide Knock-in in HEK293T cells. Precise HDR was quantified by CasAnalyzer and then filtered by NGS Analyzer to call precise insertion events

### Donorguide SpCas9 introduces insertions more efficiently than a separate ssODN donor *in vivo*

Encouraged by our *in vitro* results we sought to see if our results could be generalized to *in vivo* animal models. We chose the zebrafish model due to their high fecundity, amenability to microinjection, and ease of genotyping (Figure 1). Single cell zebrafish embryos were either injected with preformed SpCas9 RNP with a crRNA targeting the *N2B* gene in the Titin locus that we have previously shown to be highly active ^6^ (Fig 3a) along with separate “free floating” ssODN donor providing a 2nt insertion or as a CRISPR Donorguide containing the same 2nt insertion. Injected animals were incubated for 72 hours before having their genomic DNA isolated. Injected animal genomic DNA was pooled, PCR amplified and submitted for Next Generation Sequencing (Fig 1). We found that using Donorguide SpCas9, compared to WT Cas9 and a free ssODN donor, results in a similar rate of overall HDR events (Average 9.27%±4.34 for ssODN and 12.64%±2.97 for Donorguide) comprised of both precise and imprecise insertion events (Fig 3B). However, the genetic alterations induced by Donorguide SpCas9 (5.85%±1.18) resulted in a higher rate of precise insertions when compared to a free floating ssODN donor (3.44%±2.72) (Fig 3B)

**Figure 3.**
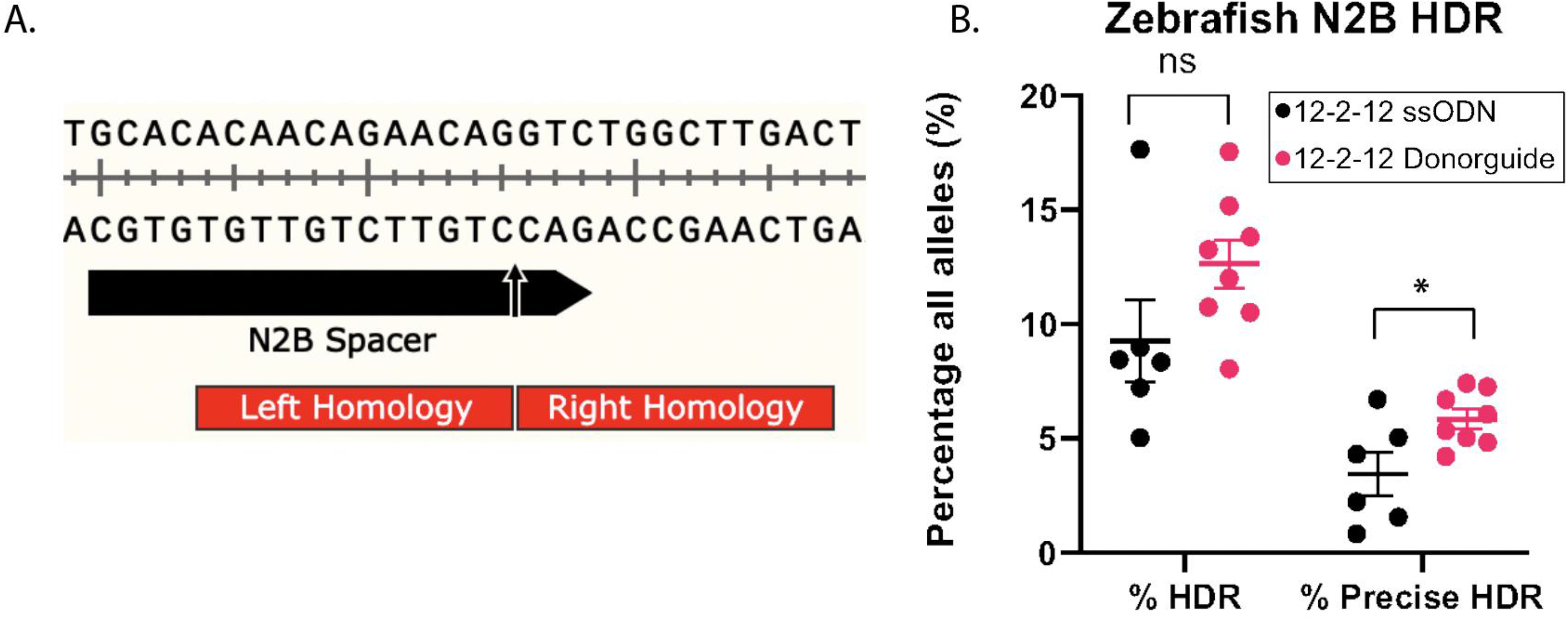
*In vivo* validation of CRISPR Donorguide at N2b. **(A)** N2b CRISPR Cas9 target site and corresponding CRISPR Donorguide Design. A 2nt insert was introduced flanked by 12nt of homolgy to the genomic target site. **(B)** Quantification of CRISPR Donorguide Knock-in in zebrafish. Precise HDR was quantified by CasAnalyzer and filtered through NGS Analyzer to call overall and precise HDR events

### Donorguide SpCas9 introduces small molecular changes more efficiently than a separate ssODN donor *in vitro*

Having shown that Donorguide is suitable to introduce small insertions *in vitro* we set out to assess if Donorguide could likewise introduce precise genetic substitutions. To this end we employed a previously established EGFP-BFP reporter system ^29^ to fluorometrically measure genetic substitution. EGFP contains 2 amino acids in its chromophore domain Thr65 and Tyr66, that when mutated to Ser65 and His66 converts it to BFP, which can be recorded by flow cytometry (Fig 4a). HEK293T cells were transfected with an EGFP expressing plasmid that is flanked by Tol2 inverted terminal repeats (ITRs) as well as a plasmid encoding tol2 transposase. Cells were passaged several times and were sorted for the brightest EGFP expression cells to establish the stable reporter cell line, termed HEK-GFP (Supplemental Figure 2).

**Figure 4.**
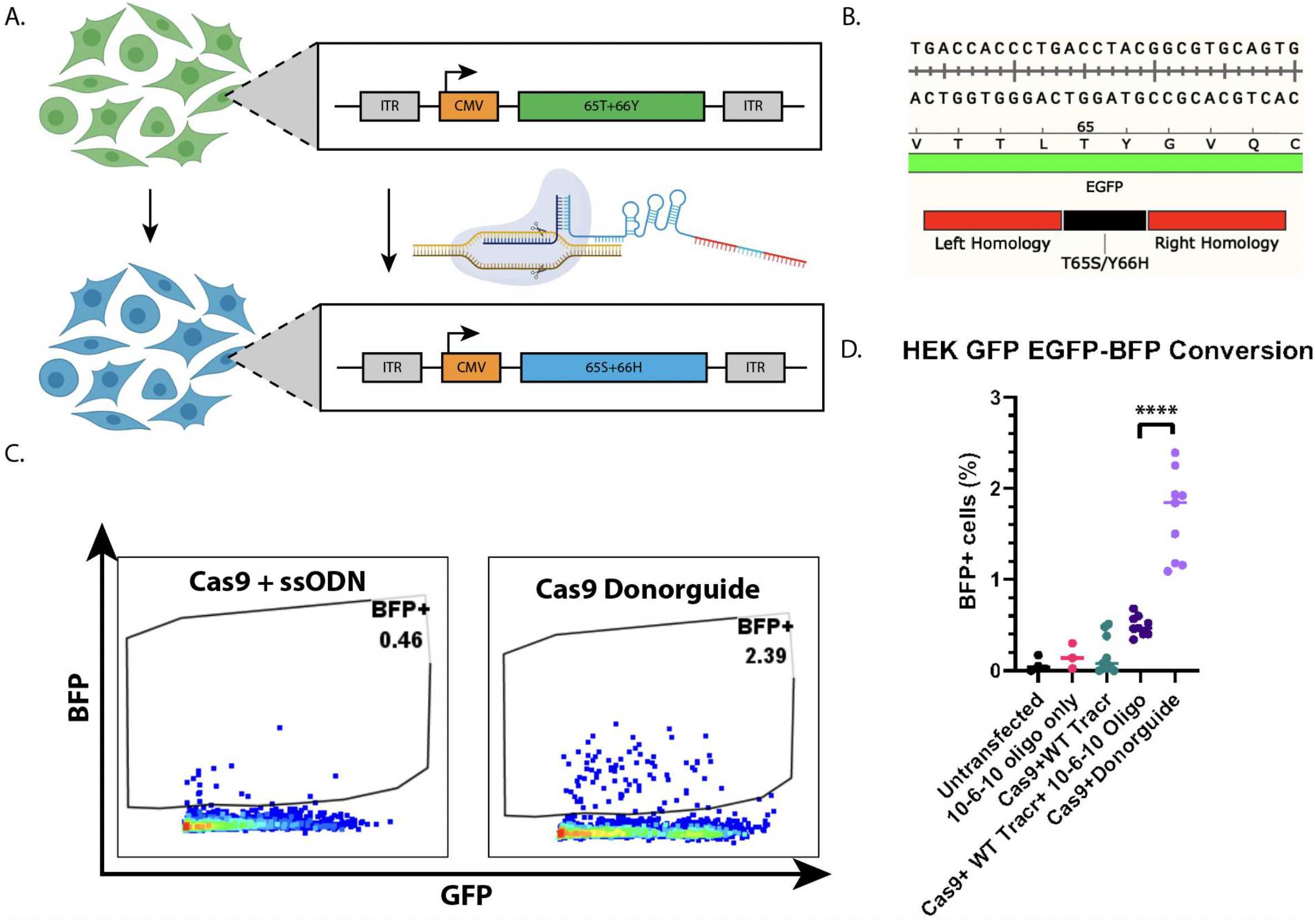
CRISPR Donorguide Converts EGFP-BFP more efficiently than an ssODN *in vitro*. **(A)** Schematic diagram of EGFP-BFP conversion cell reporter. HEK GFP cells were made by tol 2 transposition followed by FACS sorting. HEK EGFP cells were electroporated with CRISPR Donorguide or an ssODN encoding the 2 amino acid substitution flanked by 10nt of homology. Transfected HEK GFP cells were analyzed for BFP expression by flow cytometry 72 hours post transfection **(B)** Donorguide and ssODN design containing the 2 amino acid substitution flanked by 10nt of homology **(C)** Representative Flow cytometry data of HEK EGFP cells transfected with either WT SpCas9 and an ssODN encoding the EGFP-BFP substitution or with CRISPR Donorguide carrying the EGFP-BFP mutation **(D)** Quantification of BFP+ cells transfected with CRISPR Donorguide or WT SpCas9 and an ssODN donor by flow cytometry

HEK-GFP cells were transfected with Donorguide or a standard guide with a “free floating” ssODN donor along with SpCas9. The donors supplied nucleotide substitutions to produce the 2 amino acid change to convert EGFP to BFP, and this region was flanked by 10nt of homology on either side (Fig 4B). Cells were incubated at 37°C for 72 hours following electroporation and BFP expression was measured by flow cytometry. Compared to the free floating ssODN donor and WT Cas9 (average 0.49%±0.11 BFP+), Donorguide converted EGFP to BFP at a significantly greater rate (average ~1.7%±0.48 BFP+) as measured by flow cytometry (Fig 4 C-D) and next generation sequencing (Supplemental Figure 3). These results suggest that when compared to a free ssODN of similar size, CRISPR Donorguide results in a quantitative improvement of HDR induction *in vitro*.

### Donorguide SpCas9 introduces small molecular substitutions more efficiently than a separate ssODN donor *in vivo*

To assess if the conversion of EGFP-BFP observed in our HEK-GFP reporter system translated *in vivo,* we utilized a previously established transgenic zebrafish line that has an EGFP reporter stably integrated into its genome ^30^ (Fig 5A). Single cell zebrafish embryos were injected with either WT SpCas9 with an ssODN encoding the EGFP-BFP transition or CRISPR Donorguide carrying the transition. The injected animals were pooled, had their DNA isolated, barcoded, and submitted for Next Generation Sequencing. We found that compared to a free ssODN, CRISPR Donorguide (1.68%±1.25) results in more precise insertion events compared to an ssODN of a similar size (0.13%±0.17) (Fig 5B). These results suggest that CRISPR Donorguide functions cross-model to enhance HDR relative to an ssODN.

**Figure 5.**
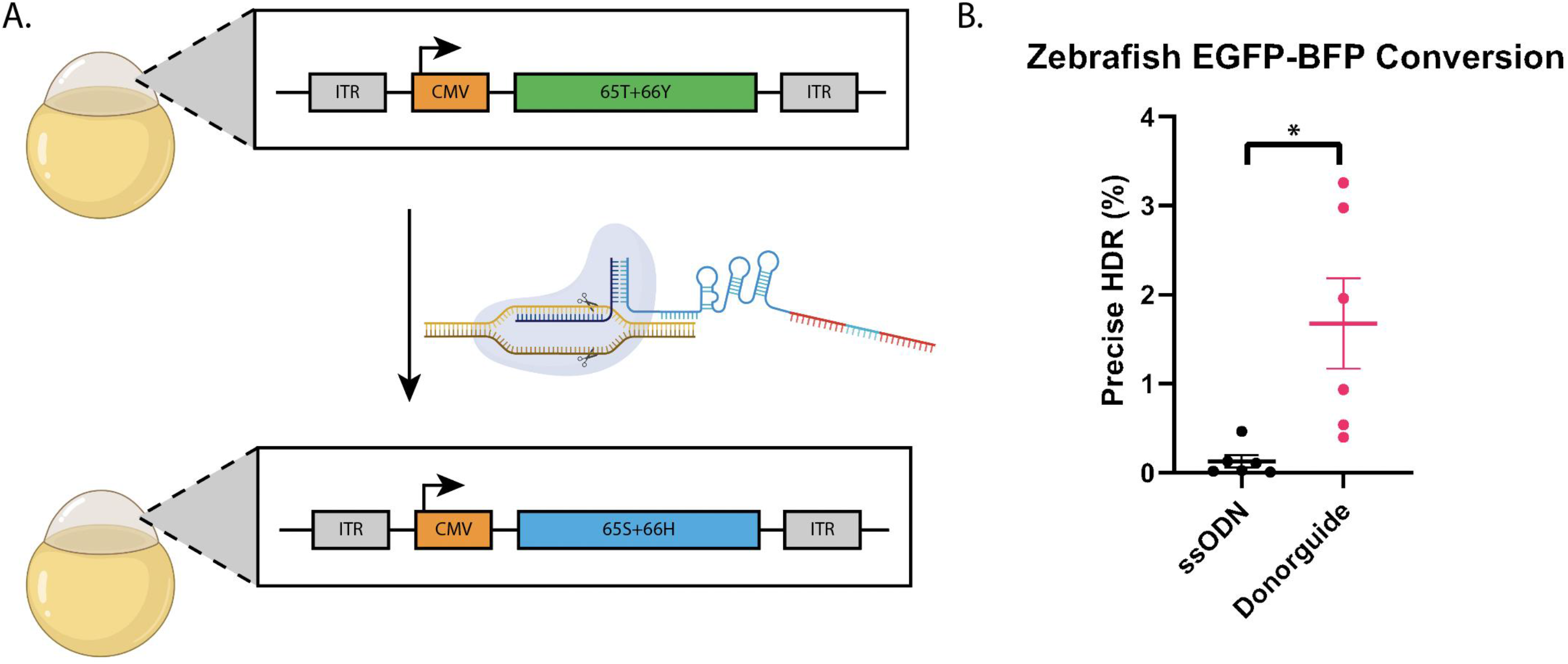
CRISPR Donorguide Converts EGFP-BFP more efficiently than an ssODN *in vivo*. **(A)** Schematic diagram of EGFP-BFP conversion in zebrafish. Transgenic zebrafish with EGFP integrated into the genome by tol2 transposition were injected at the single cell stage with either CRISPR Donorguide or WT SpCas9 and an ssODN encoding the EGFP-BFP substitution **(B)** Quantification of EGFP-BFP substitution *in vivo.* Injected animals were pooled and submitted for next generation sequencing and analyzed by CasAnalyzer and further analyzed by NGSAnalyzer to call precise or imprecise HDR events

### Donorguide SpCas9 introduces insertions more efficiently than a separate ssODN donor *in vitro*

Encouraged by our *in vivo* results, we sought to determine if the size of the homology arms explained the modest rates of HDR with CRISPR Donorguide *in vitro* (Fig 2B). To address this, we increased the homology arm length to 23nt for both CRISPR Donorguide and its requisite ssODN designed to introduce a 2nt insertion at *AAVS1* (Fig 5A). The larger 23-2-23 Donorguide or WT SpCas9 and ssODN were electroporated into HEK293T cells, incubated for 72 hours, and analyzed for HDR by Next Generation Sequencing (Fig 5a). Strikingly, we found that using larger homologies flanking the insertion resulted in precise HDR rates approaching 30% with CRISPR Donorguide and these rates were significantly greater than Cas9 and a standard ssODN donor that resulted in approximately 10% precise HDR (Fig 5b). These data suggest that CRISPR Donorguide can introduce HDR at appreciable rates and outperform a standard ssODN DNA donor.

### Donorguide SpCas9 introduces a greater proportion of desired mutant alleles than a separate ssODN donor *in vitro*

Small insertions comprise about 3% of known human pathogenic genetic variants ^27^ and being able to correct these aberrant sequences to restore the wild type gene is highly sought after. Targeting these insertion regions with CRISPR/Cas9 to resolve the lesion can result in additional loss of function alleles as a result of NHEJ or Alt-EJ repair. Therefore, precise deletions are required to address such conditions.

To this end, we wanted to determine if CRISPR Donorguide could introduce precise deletions at higher rates than a comparable ssODN. To assess precise deletion, we developed an EGFP-STOP-mRFP reporter cell line such that unperturbed cells will be EGFP+, but if a precise deletion of the stop codon is introduced the cells will be EGFP+/mRFP+ (Fig 7a). EGFP-STOP-mRFP cells were transfected with either SpCas9 and an ssODN encoding the 3nt stop codon deletion or with CRISPR Donorguide likewise containing the deletion, incubated for 72 hours, and assessed the induction of the deletion by NGS. We found that CRISPR Donorguide introduced the precise deletion at nearly the same rate as a traditional ssODN (11.45%±3.45 ssODN and 10.55%±4.03 Donorguide). However, we found that, in agreement with previous observations (Figure 2B) that Donorguide resulted in a significantly decreased amount of overall indel mutations compared to SpCas9 (88.77%±9.80 ssODN 41.23%±14.61 Donorguide). Accordingly, CRISPR Donorguide resulted in a significantly greater percentage of precise deletion alleles, defined as the number of desired alterations/overall indel mutations (25.38%±1.73 Donorguide 12.73±2.95 ssODN). These results suggest that CRISPR Donorguide can introduce a precise deletion at a comparable rate to an ssODN, but the overall fraction of desired alleles is significantly greater in the mutant allele population.

**Figure 6.**
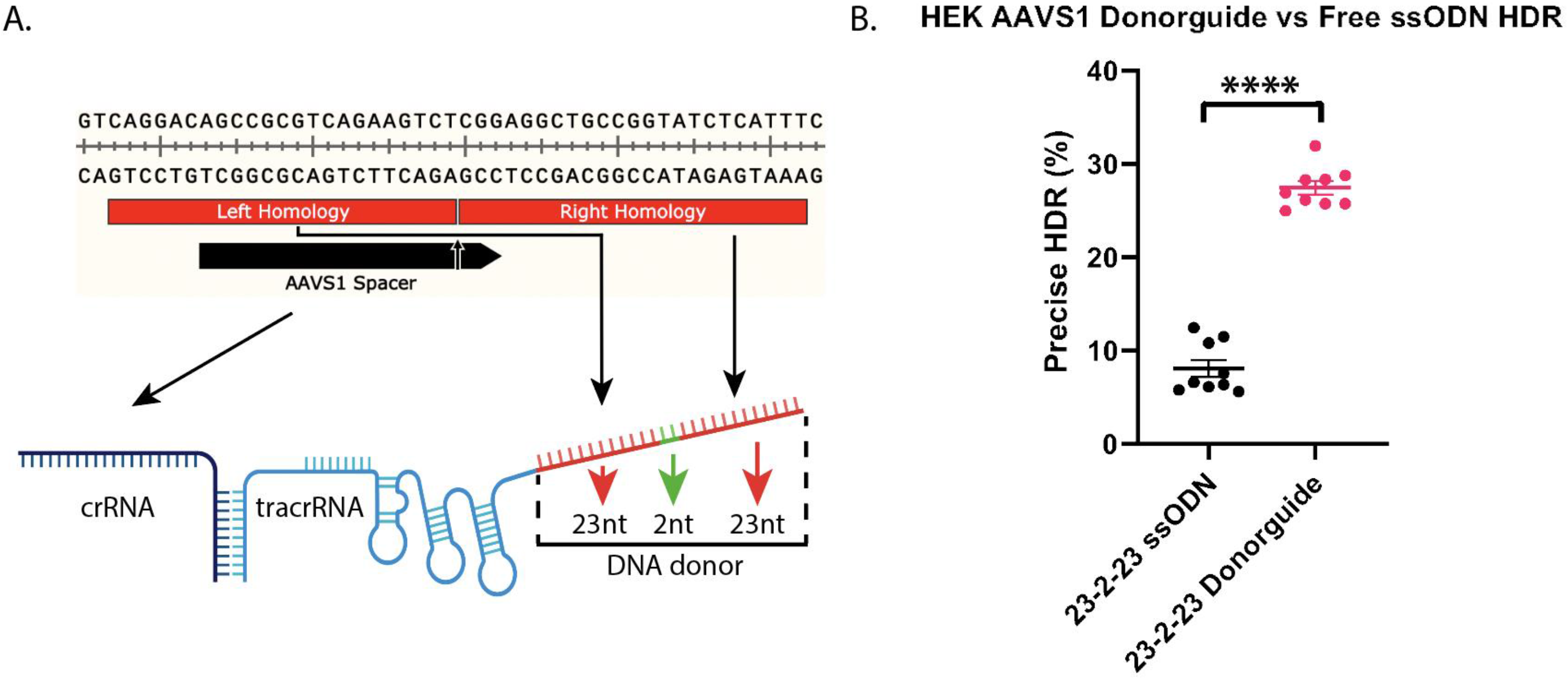
CRISPR Donorguide introduces a 2bp insertion more efficiently than an ssODN *in vitro*. **(A)** A 2nt insertion flanked by 23nt of homology was introduced by CRISPR Donorguide or with WT SpCas9 and a similar ssODN **(B)** Quantification of CRISPR Donorguide at AAVS1. Next generation sequencing data of cells transfected with either CRISPR Donorguide or WT SpCas9 and an ssODN were anaylzed by CasAnalyzer and further filtered by NGSAnalyzer to call precise insertion events

**Figure 7.**
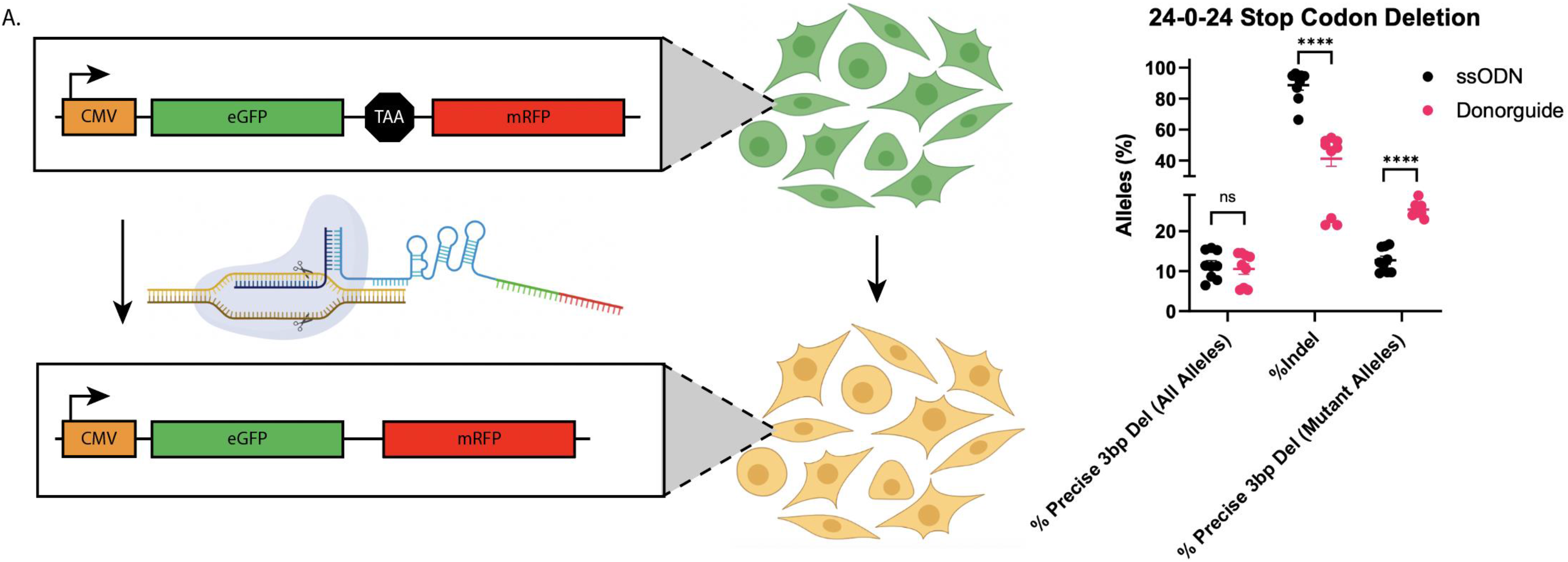
CRISPR Donorguide introduces precise deletions more efficiently than an ssODN *in vitro*. **(A)** Schematic representation of EGFP/mRFP reporter cells line. EGFP and mRFP separated by a stop codon were introduced into HEK293T cells by Tol2 transposon and sorted for the highest expressing GFP+ cells. When a donor is introduced that removes the stop codon HEKGFP cells become EGFP/mRFP double positive and can be recorded by flow cytometry. **(B)** Quantification of precise deletion in HEK GFP. HEK GFP cells transfected with CRISPR Donorguide or an ssODN encoding the 3nt deletion were analyzed for the precise 3bp deletion 72 hours post transfection

### Donorguide SpCas9 introduces a greater proportion of desired mutant alleles than a separate ssODN donor *in vivo*

As Donorguide generated a greater proportion of mutant alleles *in vitro*, we tested whether a similar phenomenon was also applicable *in vivo*. We developed an assay to effectively remove WT alleles from the population while simultaneously interrogating whether the remaining mutant alleles were generated by stochastic NHEJ-mediated indels or by a precise molecular alteration. To this end we targeted the tyrosinase (*tyr*) gene in zebrafish, which is required for converting tyrosine into the pigment melanin that confers skin color ^31^. We inserted a 6nt AvrII restriction site with either an ssODN or CRISPR Donorguide into the *tyr* locus (Figure 9A). If precisely inserted, the cargo will generate an in-frame stop codon and should lead to the development of albino zebrafish mutants (Figure 9B). The resulting albino zebrafish from both conditions were then pooled, had their genomic DNA isolated, and were PCR amplified. The resulting PCR amplicons were sub-cloned into a TA cloning vector and successfully generated TA clones were identified by blue-white screening. The resulting subclones from both the ssODN and Donorguide conditions were subjected to colony PCR followed by restriction fragment length polymorphism (RFLP) analysis using AvrII followed by agarose gel electrophoresis to visualize successfully digested (and therefore successfully integrated) alleles (Figure 9G).

**Figure 8.**
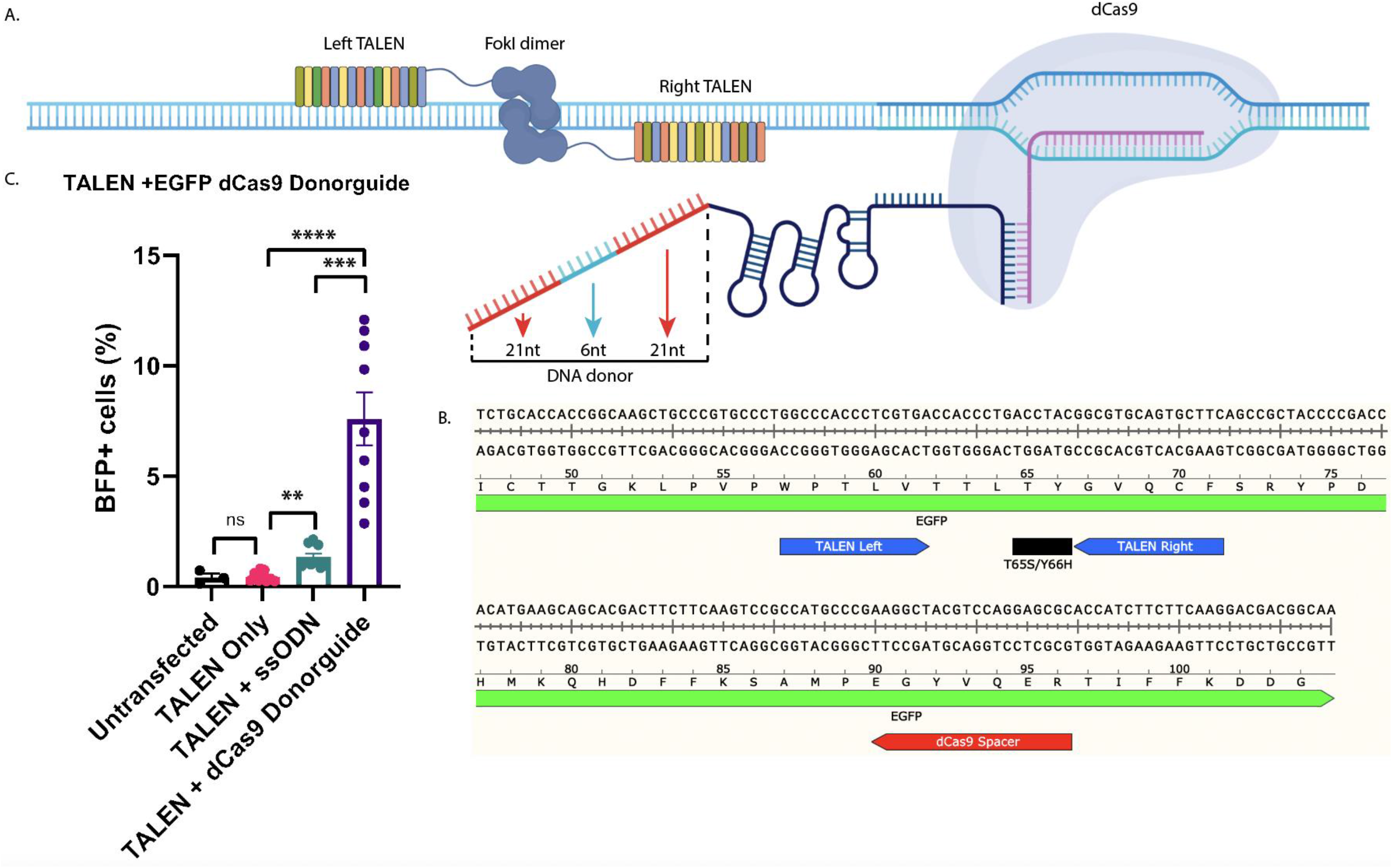
CRISPR Donorguide Enhances the Activity of other gene editing tools *in vitro*. **(A)** Schematic diagram of TALENs targeting EGFP coupled with dCas9 Donorguide. **(B)** DCas9 Donorguide targets the EGFP locus ~90nt upstream of the TALEN target site and encodes the EGFP-BFP transition flanked by 21nt of homology. **(C)** Quantification of EGFP-BFP conversion by flow cytometry. HEK GFP cells were transfected with TALENS and an ssODN encoding the EGFP-BFP transition or with TALENS and dCas9 Donorguide encoding the EGFP-BFP mutation. Transfected cells were measured for BFP expression 72 hours post transfection.

**Figure 9.**
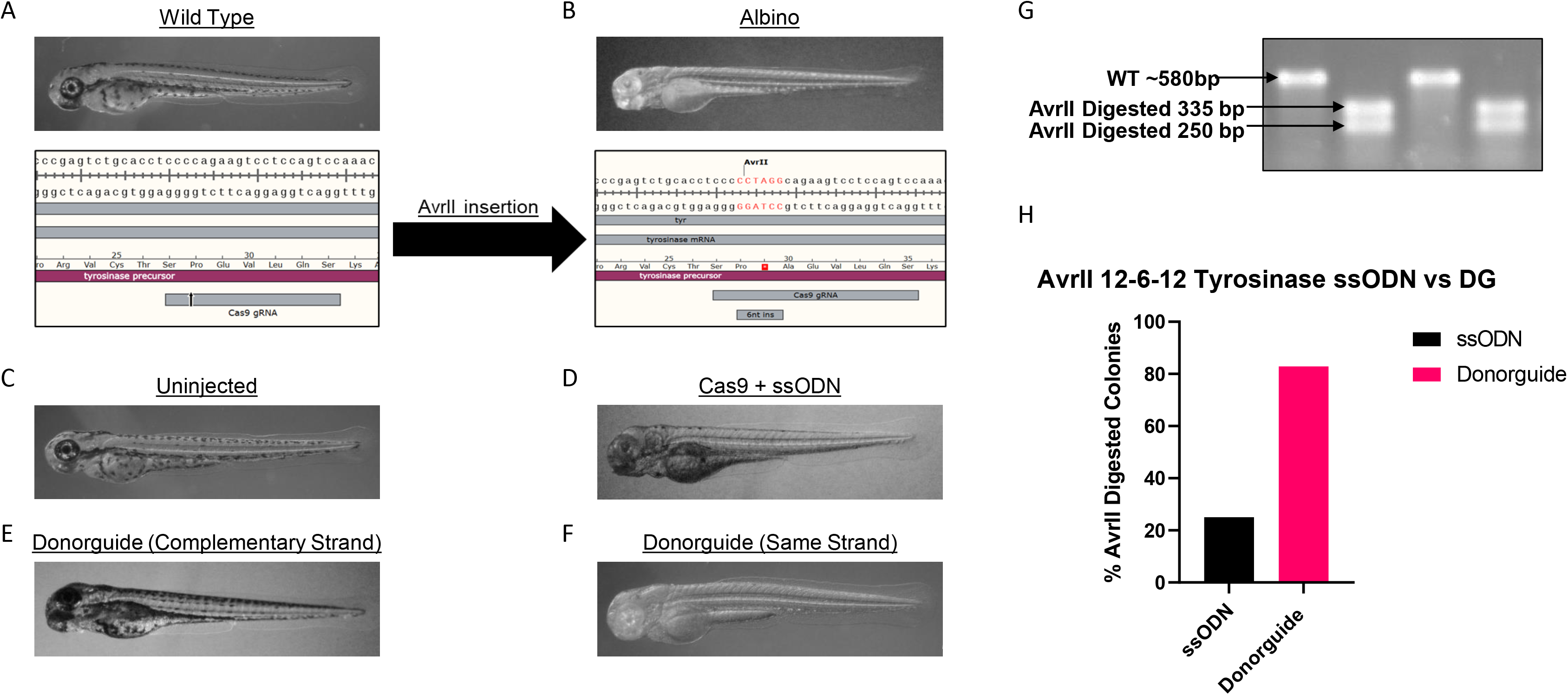
CRISPR Donorguide introduces an AvrII Site at a higher rate than an ssODN *in vivo*. **(A)** Schematic representation of the *tyr* locus and Cas9 sgRNA target site. WT *tyr* zebrafish are pigmented throughout the body and lens **(B)** Schematic representation of the *tyr* locus with the AvrII site inserted and generating a premature stop codon. The subsequent nonsense mutation results in albino zebrafish lacking body and lens pigmentation **(C-F)** Representative images of zebrafish targeted at the *tyr* locus with either an ssODN or Donorguide **(G)** Representative image of the AvrII RFLP assay showing the WT amplicon is approximately 580 but when the AvrII site is introduced and facilitates digestion, 2 bands of 335 and 250bp are generated. **(H)** Quantification on RFLP activity using either an ssODN or CRISPR Donorguide to introduce the AvrII site in *tyr*

Interestingly, while Cas9 and the ssODN donor generated albino mutants (Figure 9D) we observed only wild type looking zebrafish in the Donorguide condition (Figure 9E). Upon further inspection, we found that the DNA portion of the *tyr* Donorguide was designed complementary to the crRNA target strand. We reasoned that the DNA region of the Donorguide and the crRNA were aberrantly annealing during Donorguide assembly and preventing the formation of the crRNA:Donorguide duplex and therefore the subsequent RNP complex. We then designed the *tyr* Donorguide DNA region to be of the same strand as the crRNA target strand, and therefore prevent aberrant crRNA:Donorguide base pairing. Upon switching the orientation of the DNA donor portion, we observed strong albino phenotypes using the same strand Donorguide (Figure 9F). After performing the RFLP analysis we found that just 25% of the ssODN clones (10/40) were successfully digested with AvrII. However, we found that over 80% of the Donorguide clones (34/41) were successfully digested with AvrII. These results suggest that when eliminating WT alleles by phenotypic screening, CRISPR Donorguide comprises a majority of the mutant alleles, whereas SpCas9 and an ssODN generate mostly stochastic indel mutations.

### Donorguide works in concert with other gene editing technologies

Even with the advent of CRISPR orthologous such as Cas12a ^32^ and enhanced CRISPR Cas9 variants with decreased PAM stringency ^33^, there is still a PAM and gRNA requirement that could preclude the utilization of CRISPR tools. This is particularly an issue when trying to perform HDR, where it has been demonstrated that gene conversion is greatly enhanced when the genetic change is introduced more proximally to the site of the DNA DSB ^34^. TALENs (transcription activation like effector nucleases) on the other hand, have only a 5’ thymidine requirement, which can also be removed with mutagenic variants ^35 36^. TALENs have been shown to function efficiently as gene editing tools *in vivo ^37^ ^38^* and *in vitro ^39^* for both gene knockout and knock-in ^40^.

To this end, we wanted to assess if Donorguide could enhance TALEN-mediated HDR. We designed TALENs targeting EGFP using our previously established FusX assembly protocol (Supplemental Table 1) ^41^. We then designed a catalytically inactive Cas9 (dCas9) Donorguide targeting the EGFP chromophore domain ~80bp downstream of the TALEN target site to prevent steric interference between dCas9 and the right TALEN arm. HEK-GFP cells were transfected with both the EGFP-TALENs and an ssODN or with dCas9 Donorguide containing the EGFP-BFP transition (Fig 7a) and measured for BFP+ cells 72 hours post transfection by flow cytometry. Strikingly, we found that TALENs with dCas9 Donorguide resulted in an approximate 3-fold increase in BFP+ cells compared to TALENs with an ssODN donor. Taken together, these data suggest that CRISPR Donorguide results in greater gene conversion than a traditional ssODN donor and can be used in concert with non-Cas9 gene editing tools such as TALENs.

## Discussion

In this work we report a facile approach to introduce small genetic changes *in vitro* in HEK293T cells as well as *in vivo* in zebrafish. These results suggest that this methodology can be rapidly adapted to other cell types of interest to study protein function, validate variants of uncertain significance (VUSs), or generate isogenic cell populations to assess the implications of a patient mutation. We further show that this approach is amenable to *in vivo* editing in the zebrafish model. The success of this approach in zebrafish further suggests that this technology can be readily translated to other *in vivo* models such as mice, rats, and larger animal models.

Surprisingly we found both a qualitative and quantitative improvement over standard ssODN methods using our chimeric guide approach. In zebrafish, our data suggest that for small genetic modifications it would be preferable to use the Donorguide approach to increase both the overall amount and precision of gene integration events. This result enables researchers to better make animal models and should require less screening for allelic alterations. Imprecision of insertion events *in vivo* is well documented in the field ^42^ ^43^ ^44^ and has been partially explained by several phenomenon, but a unifying explanation has been largely elusive. We observe genetic imprecision in zebrafish in agreement with previous reports of using ssODN donors ^44^. We find that using Donorguide at least partially ameliorates some of the imprecision observed with ssODN donors. This could be explained, at least in part, by one of two factors. First, the nature of the DNA-RNA hybrid of the Donorguide could be less accessible to cellular nucleases and reduce the presence of donor fragments aberrantly integrating into the genomic lesion. Second, the Donorguide could be physically protected by the Cas9 RNP complex likewise sparing it from cellular exonucleases that ssODNs are otherwise susceptible to similarly reducing the presence of donor DNA fragments.

Conversely, there are limited reports of such imprecision in mammalian systems. Accordingly, we observe little imprecision in our *in vitro* assays. We find that in our *in vitro* studies the improvement of HDR with Donorguide compared to an ssODN is primarily an overall increase in desired mutant alleles without a significant change in precision of the events. We note one exception to this net increase of HDR events with Donorguide in our small deletion study. While we find that the net amount of precise insertion events is comparable between the ssODN and Donorguide, our approach results in a greater proportion of desired alterations in the pool of mutant alleles *in vitro*.

We recapitulated this increased mutant allele proportion phenomena at the *tyr* locus in zebrafish and showed significantly more insertion events in the Donorguide condition compared to the ssODN condition. It is worth mentioning that though there were significantly more AvrII digested clones using Donorguide (82% vs. 25% with ssODN), many of the PCR products were not completely digested (Supplemental Figure 4), suggesting that they may not be precise insertion events. Indeed, when we performed Sanger sequencing on several ssODN and Donorguide digested subclones we found a mix of both precise and imprecise insertion events.

Being able to narrow down the editing outcomes to either a desired alteration or no editing at all may be desirable in conditions such as Sickle Cell Disease (SCD) in which a point mutation causes the formation of sickled hemoglobin ^45^. Though the sickled hemoglobin is causal for the pathology of SCD, total loss of function introduced to correct the point mutation would likely be more harmful than the original point mutant. In SCD and similar pathologies, Donorguide would therefore be a preferred method of correction in which the genetic outcome is more likely to be no editing at all or a precise alteration with less collateral genetic damage.

In our *in vitro* studies we found that increasing the lengths of homology flanking the insertion resulted in a marked increase in HDR events. At this time, we are generating these chimeric tracrRNA molecules from commercial manufacturers. While it is of interest to systematically increase the length of the DNA portion of Donorguide, chemical synthesis has been a limiting factor. Current synthesis limits only allow for ~50nt of DNA be added to the 3’ end of the tracrRNA, currently limiting the possibility of cargos and lengths of flanking homology. In-house synthesis of these chimeric tracrRNA molecules could afford the addition of larger homology arms flanking diverse cargos such as peptide epitope tags or fluorescent reporter genes.

In this study, all the DNA portions of the Donorguide molecules are of the same strand as the Cas9 crRNA. Indeed, we find when we use a crRNA that is complimentary to the repair portion of Donorguide, we observe no editing *in vivo* (Figure 9E). Similarly, in our *in vitro* studies when using a Donorguide that is complementary to the crRNA, we observe the introduction of indel mutations but no HDR events (data not shown). This phenomenon can likely be explained by the crRNA aberrantly base pairing with the DNA portion of the Donorguide and preventing either the formation of the crRNA:Donorguide duplex or with the formation of the Donorguide RNP complex. We therefore developed several guidelines for those wishing to use CRISPR Donorguide in their model system: (i) design the DNA repair template of the Donorguide to be of the same strand, not complimentary to, the Cas9 crRNA (ii) For zebrafish, 12nt of homology flanking a cargo of 2-6nt should be a starting point for Donorguide design. (iii) For *in vitro* studies, designing the flanking homologies to be as large as synthesis limits allows should be the starting point.

This work provides with a streamlined technology to introduce small molecular alterations both *in vitro and in vivo* with only 2 components required. We show that CRISPR Donorguide results in either an overall increase of HDR events or an increased proportion of desired mutant alleles. Different research applications could therefore leverage Donorguide for its desired editing outcomes. We likewise include design guidelines to deploy CRISPR Donorguide in their respective model systems for enhanced precision genome editing applications.

## Methods

### Cell culture

HEK293T cells were obtained from ATCC (CRL-3216). Cells were maintained in Dulbecco’s modified Eagle’s medium (Gibco #11995-040) supplemented with 10% fetal bovine serum (Gibco #26140079) and 1% penicillin/streptomycin (Gibco #15140-122).

### *In vitro* Donorguide RNP preparation

Protocol adapted from iPS cell electroporation-NepaGene. Briefly, Alt-R CRISPR-Cas9 crRNAs were ordered from IDT (www.idtdna.com/CRISPR-Cas9) and suspended at a concentration of 200μM in nuclease free water. The chimeric CRISPR-Cas9 tracrRNAs were ordered from IDT as RNA Ultramers (https://www.idtdna.com/site/order/oligoentry/index/UltraRNA) with the general sequence “rArGrCrArUrArGrCrArArGrUrUrArArArArUrArArGrGrCrUrArGrUrCrCrGrUrUrArUrCrArArCrUrUrGrArArAr ArArGrUrGrGrCrArCrCrGrArGrUrCrGrGrUrGrCrUrUrUrUrUrUrUrUNNNNNNNNNNNNNNNNNNNN” (where rN=RNA bases comprising the tracrRNA sequence and N=DNA bases comprising the DNA donor portion) and suspended at a concentration of 200μM in nuclease free water. The specific Donorguide sequences used in this study can be found in Supplemental Table 1.To assemble the Donorguide for *in vitro* studies, 3ul of 200μM crRNA was combined with 3μl of 200μM chimeric tracrRNA and heated to 95°C for 5 minutes and immediately removed to incubate on the bench top at RT for 10-20 minutes until crRNA:tracrRNA mix reached RT. 5μl of the 100μM crRNA:tracrRNA mix was added to a new nuclease free PCR tube. 4μl of Alt-R Cas9 Nuclease 3NLS (62 μM stock) (IDT cat # 1074181 1074182) was added to bring to a final volume of 9μl and incubated at RT for 20-30 minutes to form the Donorguide RNP complex.

### Transfections

For Donorguide HEK293T electroporations, 9ul of the preformed Donorguide RNP complex (see above) was diluted with 16μl in EP buffer (Etta Biotech) to bring to a final volume of 25μl. Diluted Donorguide RNP complex was added to 300μl HEK293T cells in EP buffer at a concentration of 20E6cells/ml, mixed thoroughly, and 107μl of the HEK293T cell+ Donorguide cocktail was added to each cuvette.

For the TALEN dCas9 Donorguide electroporations, 9μl of the preformed dCas9 Donorguide RNP cocktail and 5μl of 100μM ssODN with 4μl PBS was brought to 25μl with 16μl of EP buffer. Diluted Donorguide RNP complex or ssODN mix was added to 300μl HEK293T cells in EP buffer at a concentration of 20E6cells/ml, mixed thoroughly, and 107μl of the HEK293T cell+ Donorguide cocktail was equally distributed to 3 Eppendorf tubes. 2.5μg of each left and right TALEN arm plasmids were added to each tube and then transferred to an electroporation cuvette.

HEK293T cells were then electroporated with Etta H1 electroporator (Etta Biotech) with the following parameters: 200 V, 784 ms interval, six pulses, and 1,000 μs pulse duration. Cells are recovered post electroporation by incubating at 37°C for 5–10 min before being transferred to a six-well tissue cell culture plate at a density of about 1.5E6 cells/mL.

### DNA isolation and PCR Amplification

DNA from whole-cell populations was purified using the Qiagen DNeasy Blood & Tissue Kit (Qiagen 69504). DNA was isolated from zebrafish as described below. PCR amplification was conducted with Platinum SuperFi DNA Polymerase (Thermo Fisher Scientific 12351010) or Q5® High-Fidelity 2X Master Mix (New England Biolabs M0492S) and purified with the Qiagen QIAquick PCR Purification Kit (Qiagen 28104).

Samples used for RFLP analysis were amplified with MyTaq™ Red Mix (Bioline BIO-25043) and subcloned using Strataclone PCR Cloning Kit (Agilent 240205). Either Donorguide or ssODN injected zebrafish were combined, isolated and PCR-amplified in three separate reactions and subjected to three separate subcloning steps to represent as many alleles as possible. Colony PCR was performed with M13F/M13R using MyTaq™ Red Mix and amplification confirmed by agarose gel electrophoresis. The colony PCR amplified products were digested with AvrII (New England Biolabs R0174S) in CutSmart buffer at 37°C for 3 hours before being resolved with agarose gel electrophoresis.

### GeneWiz AmpliconEZ: DNA library preparation and Illumina sequencing

PCR products were obtained and purified as mentioned above. Illumina partial adapters (Forward sequencing read: 5’-ACACTCTTTCCCTACACGACGCTCTTCCGATCT-3’ reverse sequencing read: 5’-GACTGGAGTTCAGACGTGTGCTCTTCCGATCT-3’) were added to the 5’ end of sequencing primers. DNA library preparations, sequencing reactions, and initial bioinformatics analysis were conducted at GENEWIZ, Inc. DNA library preparation, clustering, and sequencing reagents were used throughout the process using the NEBNext Ultra DNA Library Prep Kit, following the manufacturer’s recommendations (Illumina). End-repaired adapters were ligated after adenylation of the 3′ ends followed by enrichment by limited cycle PCR. DNA libraries were validated on the Agilent TapeStation, quantified using the Qubit 2.0 Fluorometer (Invitrogen), and multiplexed in equal molar mass. The pooled DNA libraries were loaded on the Illumina instrument according to the manufacturer’s instructions. The samples were sequenced using a 2 × 250 paired-end (PE) configuration. Image analysis and base calling were conducted by the Illumina Control Software on the Illumina instrument.

### Next Generation Sequence Analysis

The raw .fasta files obtained from GeneWiz AmpliconEZ were analyzed using CasAnalyzer ^46^ to determine indel efficiency and overall HDR events. The raw output from CasAnalyzer was then further analyzed by NGS Analyzer to quantify precise and imprecise HDR events. This script for NGS Analyzer is hosted at https://github.com/srcastillo/Sequencing under a GNU General Public License v3.0.

### TALEN design and assembly

A pair of TALENs against the *EGFP* gene were designed and generatedusing the FusX assembly system ^47^. The RVDs utilized were HD = C, NN = G, NI = A, NG = T. The TALEN pair binds the following *EGFP* gene sequence, where the spacer region, which is the site of predicted DSBs, is in lower case: GGCCCACCCTCGTGAccaccctgacctacGGCGTGCAGTGCTTC. TALEN plasmids were electroporated into HEK-EGFP cells. After electroporation, substantial gene editing in the bulk population was verified.

### Zebrafish husbandry

Wild-type (WT) MRF zebrafish (*Danio rerio*) were maintained in the Mayo Clinic Zebrafish Core Facility. Fish are handled in accordance with standard practices ^48^ and guidelines from the Institutional Animal Care and Use Committee (IACUC) (A34513-13-R16, A8815-15). Adult zebrafish are kept in a 9 L (25 – 30 fish) or 3 L housing tanks (10 – 15 fish) at 28.5°C with a light/dark cycle of 14/10 hours. Adult fish pair breeding was set up a day before microinjections, and dividers were removed the following morning. After microinjections, the fertilized embryos were transferred to Petri dishes (100 mm) in E2 media (1x; 15 mM NaCl, 0.5 mM KCl, 1 mM MgSO4, 150 μM KH2PO4, 50 μM Na2HPO4, 1 mM CaCl2, 0.7 mM NaHCO3) and maintained in an incubator at 28.5°C. Sexes are not determined in larval fish (2 – 5 days post-fertilization (dpf)) in this study and thus potential differences based on sex are not assessed. Zebrafish gonads begin developing around 15 dpf ^49^. A multifactorial sexual differentiation process, involving multiple genes and environmental factors, takes place around 25 dpf and is considered completed around 60 dpf (^50–52^).

### DNA oligonucleotide preparation

All single strand oligonucleotides (ssODNs) used in this study were purchased from IDT (Integrated DNA Technology, Inc., Coralville, IA, USA). Stock was prepared at 100 μM in 1× TE and working stock was prepared at 20 μM to be used for microinjection or left at 100 μM for *in vitro* studies.

### crRNA, tracrRNA, Donorguide, and spCas9 preparation for zebrafish microinjection

All crRNA (crispr RNA), tracrRNA (transactivating crispr RNA), Donorguide (tracrRNA fused with a DNA cargo sequence flanked by homology arms), and SpCas9 (Streptococcus pyogenes Cas9) were custom-ordered from IDT. Stock was prepared at 20 μM in Duplex Buffer (IDT; Cat#11-05-01-03; 30 mM HEPES, pH 7.5, 200 mM potassium acetate) for crRNA, tracrRNA, and Donorguide. At the 20 μM concentration, crRNA is at 233.9 pg/nL (protospacer length 20 nt) and tracrRNA is at 432.7 pg/nL. spCas9 stock was prepared at 23 μM (3.73 ng/nL) in PBS (phosphate buffered saline; 1×).

### Microinjection

1-cell stage fertilized zebrafish embryos were harvested and aligned on an agarose plate in E2 media. 3 μL of injection solution was loaded into an injection needle (a broken pulled glass pipette). A 2 nL of injection solution was delivered into the cell interface layer of the embryo. This injection (2 nL) delivered crRNA:tracrRNA (or Donorguide):spCas9 complex at approximately 4.52 fmol (52.9 pg):4.52 fmol (97.8 pg):4.52 fmol (733.1 pg) and ssODN (when needed) at 4 fmol. Injected embryos were transferred to a new Petri dish and kept in an incubator at 28.5 °C. Dead and nonviable embryos were counted and removed the next day (1 dpf).

### Zebrafish experimental replicates

All experiments involving zebrafish microinjection were carried out in three independent experiments. Each experiment utilized a mixed populations of embryos obtained from multiple clutches. Each experiment was ideally conducted on different days. If two experiments were performed on a day, all reagents were prepared from different stock preparations.

### Zebrafish genomic DNA extraction

Usually 40 embryos were individually collected in 36 μL of NaOH (50 mM) on 1-2 dpf (if morphological phenotyping was not necessary) and on 4-5 dpf (if morphology was assessed). Each embryo in NaOH was heated at 95°C for 15 min while vortexing a few times (REF#6-8). To stabilize NaOH, 4 μL of tris (1 mM) was added. 8 embryos were pulled into a sample (a biological replicate) by taking 2 μL from each embryo (8 fish per sample), and a 1:3 dilution was made by adding 48 μL of distilled water.

## Author contributions

SCE, KJC, HA, and BWS conceived the study. BWS, KJC, SCE, and HBL wrote the manuscript. HA performed proof of concept zebrafish experiments. CD and HBL performed zebrafish experiments. BWS performed cell line experiments and analyzed all data sets. SRC wrote the NGS Analyzer script and assisted with figure and TALEN design. GMG designed the figures. WAG provided preliminary cell line data.

## Acknowledgements

We would like to thank the Mayo Clinic Zebrafish Facility staff for assisting with animal husbandry and the Mayo Clinic Fluorescent Microscopy core for assistance with flow cytometry. We would also like to thank Dr. Alex Abel for assistance with flow cytometry analysis. We would like to thank Sharoon Akhtar and Xiyin Wang for insightful discussion that informed the experiments. We additionally thank Dr. Maura McGrail and Dr Jeff Essner for their constructive collaboration and advice. This work was supported by NIH R24 OD020166 (SCE, KJC), Mayo Foundation for Medical Education and Research (BWS, SRC, WAG, GMG), Mayo Clinic Regenerative Sciences Training Program (BWS), and the Harry C. and Debra A. Stonecipher Predoctoral Fellowship (SRC).

## Supplemental Figure Legends

**Supplemental Figure 1.**
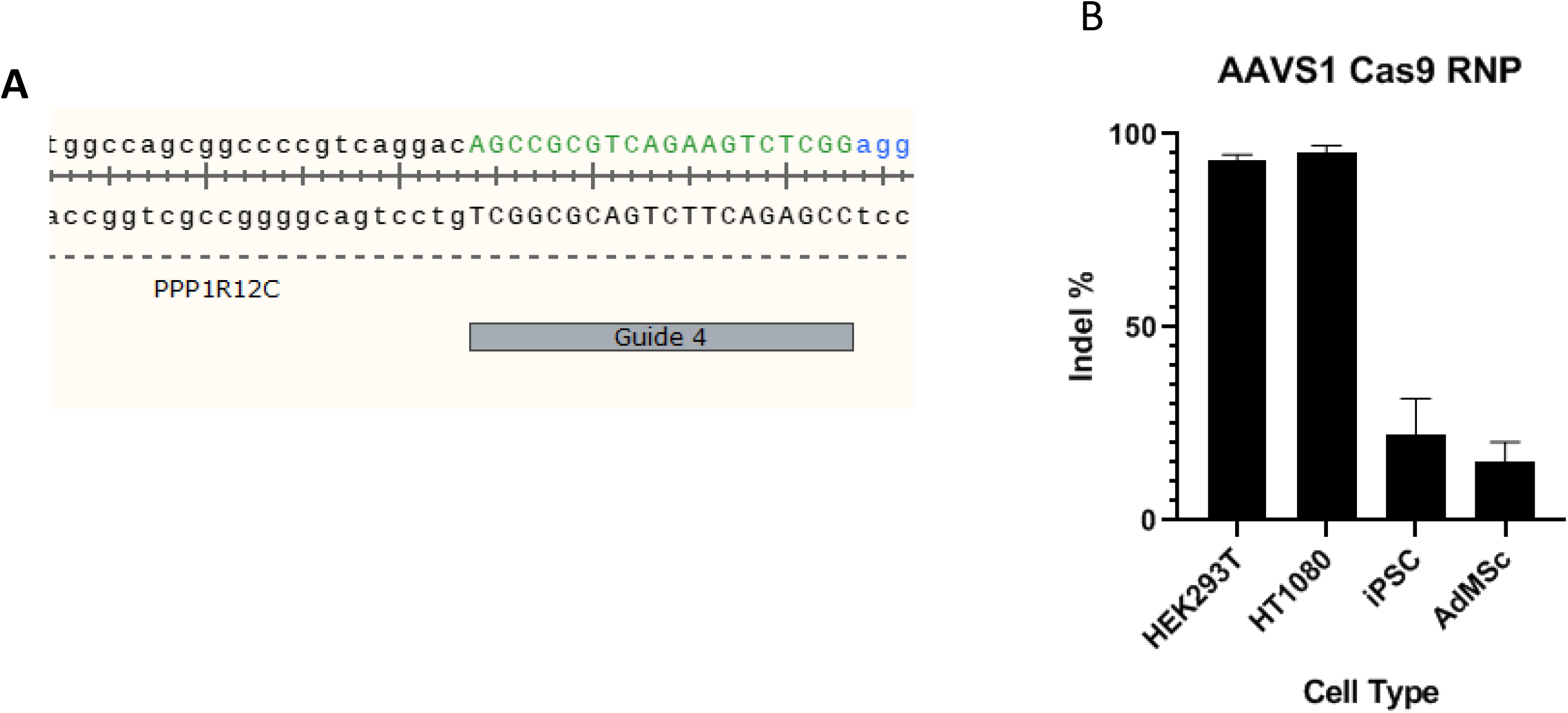
In vitro validation of SpCas9 sgRNA at *AAVS1*. (A) CRISPR/Cas9 AAVS1 genomic target site (B) Quantifcation of Cas9 activity at AAVS1 in various cell types by Sanger Sequencing and ICE Analysis

**Supplemental Figure 2.**
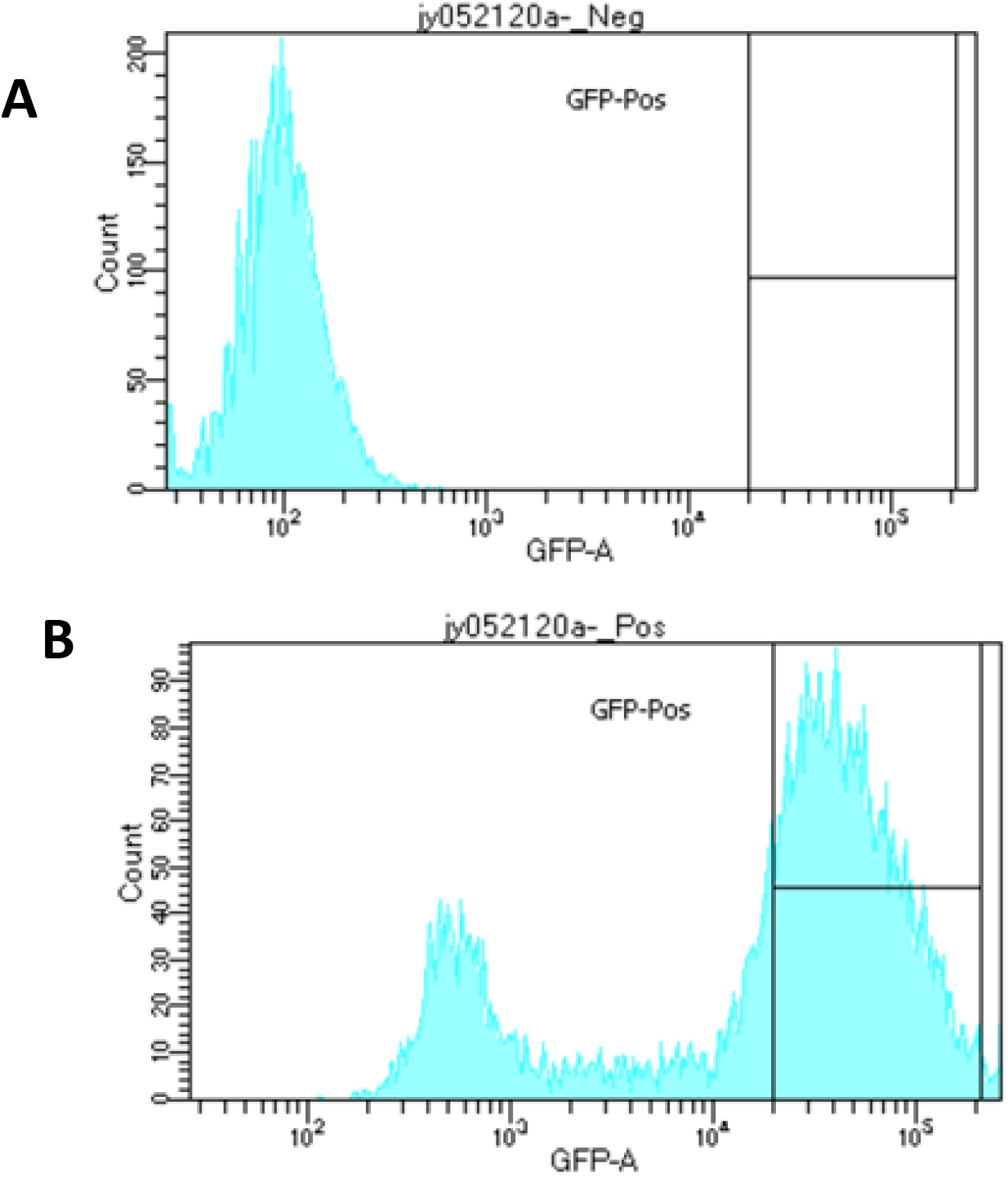
Generation of HEK EGFP Reporter Cell line. **(A)** Untransfected HEK 293T cells **(B)** HEK 293T cells transfected with pkTol2cEGFP plasmid + pkC Tol2 plasmid and passaged ~10 times. Gating indicates EGFP cells sorted to generate HEK EGFP cell line

**Supplemental Figure 3.**
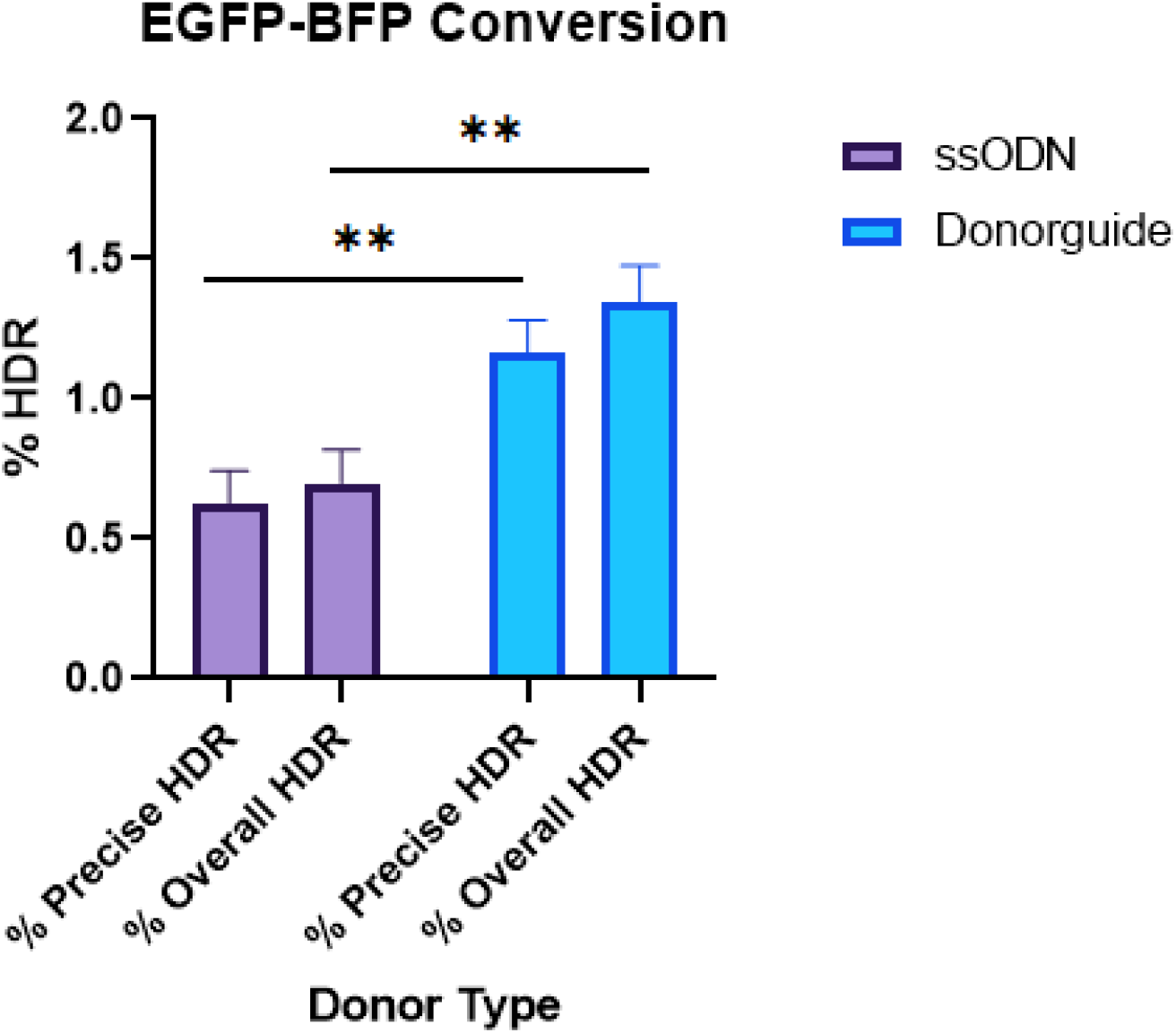
Donorguide converts EGFP-BFP *in vitro*. Next Generation Sequencing confirmation of EGFP-BFP transition with CRISPR donorguide or an ssODN in HEK GFP reporter cells.

**Supplemental Figure 4.**
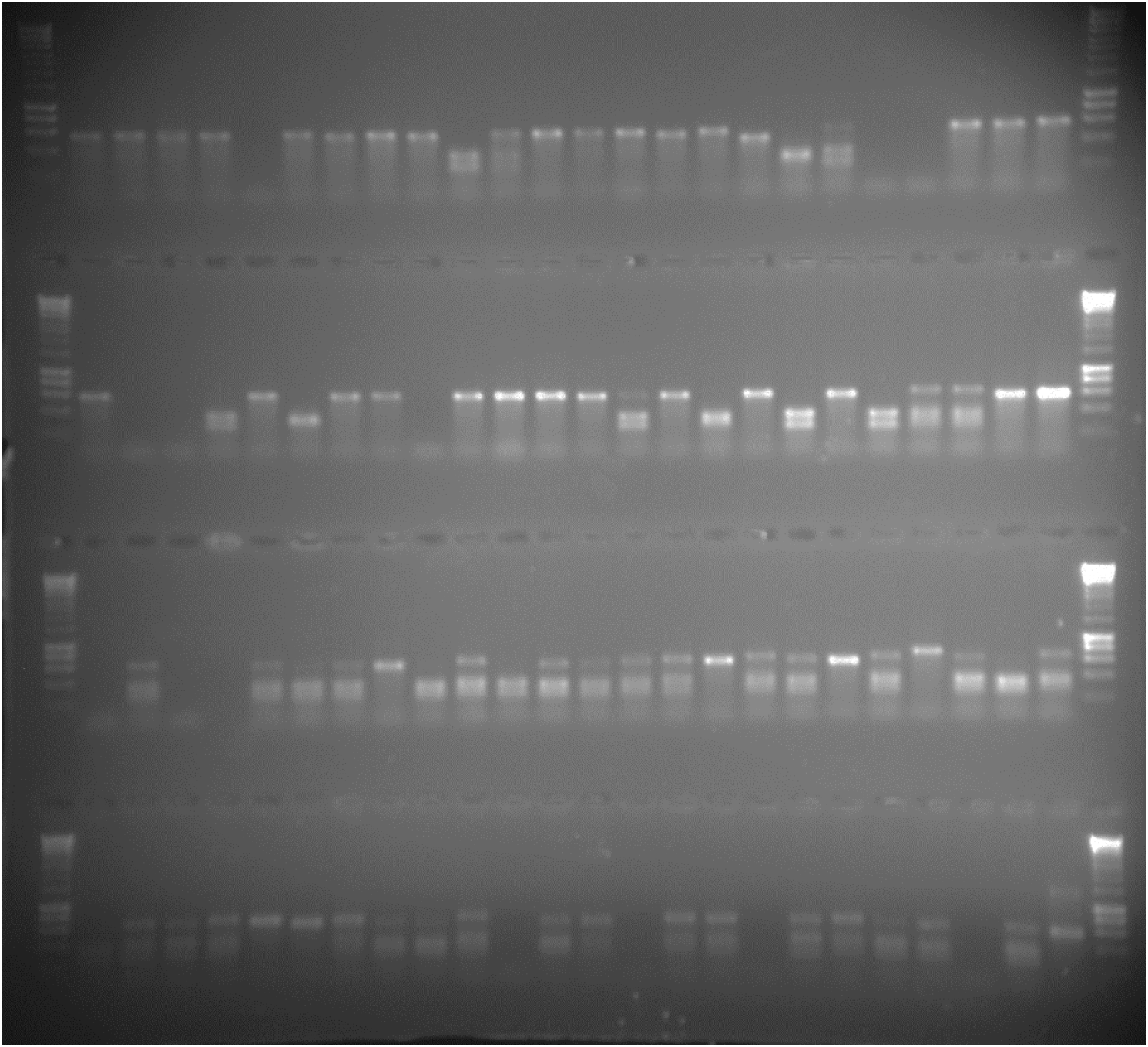
Donorguide introduces an AvrII restriction site at a higher rate than a comparable ssODN. (Top two rows) ssODN injected subclones amplified with colony PCR and digested with AvrII. 8 clones were excluded for either failing to amplify or amplifying a band the size of an empty strataclone vector. 10 clones successfully digested with AvrII (Bottom two rows) Donorguide injected subclones amplified with colony PCR and digested with AvrII. 7 clones were excluded for failing to amplify 34 clones successfully digested with AvrII.

